# Dissociation of Reward Incentive and Motivational Drive by Distinct Neural Circuits in Mice

**DOI:** 10.64898/2026.11.21.768340

**Authors:** Makoto Takemoto, Wen-Jie Song

## Abstract

The generation of reward incentive value and maintenance of motivational drive are distinct processes in reward-related behavior. However, how neural circuits differentially process these components remains poorly understood. Although intracranial self-stimulation (ICSS) paradigms have been utilized to identify reward-related circuits, ICSS measures may conflate reinitiated actions attributed to learned incentive value with ongoing motivated behavior sustained by concurrent stimulation. Here, we reveal a dissociation between these processes by optogenetically stimulating fibers in the stria terminalis bundle projecting to the medial nucleus accumbens (STB^mNAc^) and those in the medial forebrain bundle originating from the lateral NAc (MFB^lNAc^). We found that STB^mNAc^ stimulation facilitated the reinitiation of ICSS actions, whereas MFB^lNAc^ stimulation sustained ongoing ICSS behavior. Moreover, shortening the stimulation duration in STB^mNAc^ mice increased the likelihood of subsequent re-engagement of stimulation-seeking. Additionally, STB^mNAc^ stimulation, but not MFB^lNAc^ stimulation, could substitute for primary rewards and support the acquisition of sensory cue-dependent operant behavior. Together, our findings suggest that STB^mNAc^ activation promotes reward-anticipatory behavior by conferring reward incentive value, whereas MFB^lNAc^ activation sustains ongoing behavior by maintaining motivational drive. Thus, reward-related behavior may not be governed by a unitary motivational system but may involve anatomically and functionally distinct neural components.

## Introduction

Reward-related behavior comprises multiple processes, including the acquisition of reward incentive value, the anticipation of rewarding outcomes, and the initiation and persistence of reward-seeking actions (*1, 2*). An important question is how distinct neural circuits contribute to the different components of reward-related behavior.

Intracranial self-stimulation (ICSS) paradigms allow animals to repeatedly perform actions that trigger stimulation of specific brain regions or neural circuits. Because the reinforcing effects of ICSS are independent of peripheral sensory input (*3*), these paradigms have been widely used to identify the neural substrates of reward-related behavior in rodents (*3–6*). However, conventional ICSS measures do not readily distinguish between anticipatory behavior driven by previously learned reward incentive value and the persistence of ongoing behavior sustained by concurrent stimulation. Even in real-time preference paradigms, increased time spent at a stimulation-associated option, commonly interpreted as an indication that stimulation confers reward incentive value (*7–10*), may instead reflect prolonged engagement in ongoing behavior, regardless of whether reward incentive value is acquired. Moreover, under natural conditions, reward acquisition typically interrupts further seeking, at least temporarily, through consumption or transient satiety, whereas ICSS can sustain ongoing seeking behavior. This discrepancy raises the fundamental question of whether ICSS confers reward incentive value or primarily sustains ongoing behavior.

Cued operant conditioning provides a complementary approach to determine whether stimulation of neural circuits can itself generate reward incentive value and establish cue-evoked reward-anticipatory behavior (*11, 12*). Unlike repetitive responding observed during conventional ICSS, cue-dependent operant behavior can reveal whether prior stimulation experience supports the subsequent initiation of reward-seeking actions. Although primary rewards, such as food and water, readily support such learning, it remains unclear whether stimulation of specific reward-related circuits can substitute for primary rewards in establishing cue-dependent operant behavior.

Optogenetic studies have identified neuronal populations in several brain regions that support reinforcement in ICSS paradigms, including the ventral tegmental area (VTA) (*13, 14*), nucleus accumbens (NAc) (*10, 15–17*), basolateral amygdala (BLA) (*7, 18–20*), lateral hypothalamus (*21*), and medial prefrontal cortex (*22, 23*). Among these regions, anatomical studies have shown that the NAc receives projections from the BLA (*24–26*) and that axons from a population of BLA neurons course through the stria terminalis (ST) to innervate multiple forebrain regions, including the NAc (*27, 28*). Functionally, the BLA-NAc pathway has been implicated in behavioral reinforcement (*10, 17, 20, 29, 30*) and the association between primary rewards and predictive cues (*18–20, 31–34*). These findings suggest that the ST may provide an anatomical route through which the BLA contributes to reward-related behavior. However, it remains unknown whether the selective stimulation of NAc-projecting axons at the ST can provide reward incentive value and thereby establish cue-dependent reward-anticipatory behavior.

Here, we sought to dissociate reward-dependent reinitiation of actions from stimulation-driven behavioral persistence by stimulating distinct neural pathways associated with the NAc, a central hub of reward-related circuits (*3*). Using virus-mediated tracing, we identified a prominent fiber bundle coursing through the ST and projecting to the medial NAc (mNAc), which we termed the mNAc-projecting ST fiber bundle (STB^mNAc^). We investigated the behavioral effects of STB^mNAc^ stimulation compared with stimulation of the lateral NAc-derived descending axons at the medial forebrain bundle (MFB^lNAc^), a canonical substrate of ICSS (*4, 5, 35*). By combining optogenetic stimulation with behavioral analyses that separately quantify the frequency of action reinitiation and the duration of individual actions, we found that STB^mNAc^ stimulation promoted the reinitiation of seeking behavior, whereas MFB^lNAc^ stimulation predominantly sustained ongoing behavior. Moreover, STB^mNAc^ stimulation supported cue-dependent operant conditioning in the absence of primary rewards. Together, these findings suggest that STB^mNAc^ activation generates reward incentive value that promotes the reinitiation of anticipatory behavior, whereas MFB^lNAc^ activation maintains motivational drive that promotes the persistence of ongoing behavior.

## Results

### Selective labeling of an mNAc-projecting fiber bundle in the stria terminalis

First, we explored the pathway of the NAc afferents in mice to determine a suitable site for optogenetic stimulation to investigate reward-related behavior. Following injection of a retrograde adeno-associated virus (AAV) expressing a fluorescent protein into the mNAc (Fig. 1a), we observed a prominent fluorescently labeled fiber bundle coursing through the ST (Fig. 1b), which we refer to as the STB^mNAc^. In line with previous studies demonstrating that the mNAc receives input from the posterior BLA (pBLA) (*17, 31, 32*), retrogradely labeled cell bodies were detected in the BLA, with a higher density observed in its posterior half along the anterior-posterior (A-P) axis (Fig. 1b−d). The number of labeled cells was significantly greater in the pBLA than in the anterior BLA (Fig. 1d; paired t test, *P <* 0.001, *n =* 5 mice).

**Figure 1.**
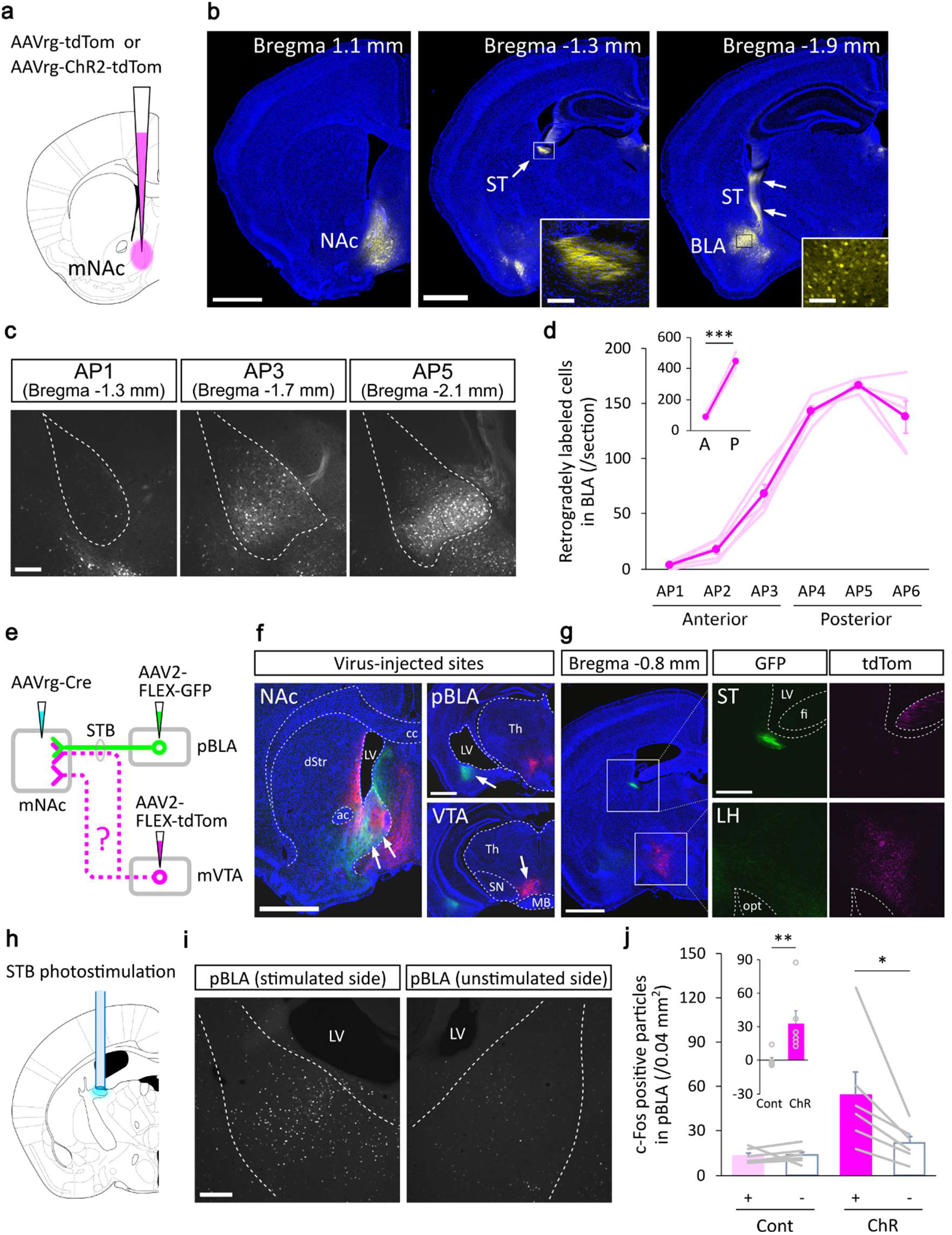
Identification of the STB pathway projecting to the mNAc. **a** Schematic representation of retrograde AAV injection into the mNAc. **b** Representative images of coronal sections showing a virus-injected site (left panel) and a tdTomato-labeled (yellow) fiber bundle in the ST (arrows in the middle and right panels). Retrogradely labeled cell bodies were observed in the BLA (right panel). Insets show confocal magnification of the boxed regions. The sections were counterstained with DAPI (blue). Scale bars: 1 mm (left and middle panels); 0.1 mm (insets). **c** Representative grayscale images showing the distribution of retrogradely tdTomato-labeled cells in the BLA at the levels of 1.3 mm (AP1), 1.7 mm (AP3), and 2.1 mm (AP5) posterior to bregma. Bregma levels were determined based on a mouse brain atlas (*45*). Scale bar: 0.2 mm. **d** Quantification of retrogradely labeled cells across the anterior-posterior axis of the BLA. AP2, AP4, and AP6 were 1.5 mm, 1.9 mm, and 2.3 mm posterior to bregma, respectively. Light colors represent individual mice, and a dark color represents the mean. The inset shows the number of retrogradely labeled cells in the anterior half (A, the sum of AP1-AP3) and posterior half (P, the sum of AP4-AP6) of the BLA (paired t test, *t*(4) = −20.68, *P <* 0.001, *n =* 5 mice). **e** Experimental strategy for the selective labeling of mNAc-projecting neurons in the pBLA and mVTA using a retrograde AAV-Cre-mediated approach. GFP and tdTomato were expressed in a Cre-dependent manner to visualize the pBLA-mNAc and mVTA-mNAc pathways. **f** Representative images of coronal sections showing virus-injected sites in the mNAc (AAVrg-Cre; arrows on the left), pBLA (AAV2-FLEX-GFP; arrow on the top right), and mVTA (AAV2-FLEX-tdTom; arrow on the bottom right). GFP-and tdTomato-labeled axons were found in the mNAc with distinct projection patterns. Scale bars: 1 mm. **g** Representative images of a coronal section obtained from the same mouse in f, showing a GFP-labeled pBLA-mNAc axon bundle coursing through the ST (green), whereas tdTomato-labeled mVTA-mNAc fibers were virtually absent in the ST but were prominent in the lateral hypothalamic area (magenta). Each of the middle (GFP) and right (tdTom) panels shows confocal images of the two boxed regions in the left panel. Scale bars: 1 mm (left); 0.4 mm (right). **h** Experimental design for optogenetic STB^mNAc^ photostimulation. ChR2 was expressed in mNAc-projecting neurons using retrograde viral delivery, as shown in a. Blue light was delivered to the ST using optical fibers. **i** Representative images of c-Fos immunoreactivity in the pBLA following unilateral STB^mNAc^ photostimulation. Scale bar: 0.2 mm. **j** Quantification of c-Fos-immunostained particles in the pBLA of the STB^mNAc^-illuminated (+) and intact (−) hemispheres. No significant interhemispheric difference was found in tdTomato-control mice (Cont; Wilcoxon signed-rank test, *V* = 6, *P =* 0.40, *n =* 6 mice), whereas the number of c-Fos^+^ particles was significantly larger on the illuminated side than on the intact side in ChR2-expressing mice (ChR; Wilcoxon signed-rank test, *V* = 21, *P =* 0.031, *n =* 6 mice). The inset shows the difference in the number of c-Fos^+^ particles between the illuminated and intact hemispheres in Cont (*n =* 6) and ChR (*n =* 6) mice (Wilcoxon rank sum test, *W* = 1, *P <* 0.01). Lines and circles indicate individual mice, and bars represent means ± SEM. \**P <* 0.05, \*\*\**P <* 0.001. Abbreviations: ac, anterior commissure; cc, corpus callosum; dStr, dorsal striatum; fi, fimbria of the hippocampus; LH, lateral hypothalamus; LV, lateral ventricle; MB, mammillary body; opt, optic tract; SN, substantia nigra; Th, thalamus.

To clarify whether the mNAc-projecting population of pBLA neurons is a source of the STB^mNAc^, we selectively labeled mNAc-projecting neurons in the pBLA with GFP, using a retrograde Cre-dependent strategy (Fig. 1e, f). For comparison, we also labeled another source of mNAc-projecting neurons located in the medial ventral tegmental area (mVTA) (*16, 17*) with tdTomato in the same brain (Fig. 1e, f). GFP-labeled axons originating from the pBLA were observed as a dense fiber bundle within the ST, whereas mVTA-derived tdTomato-labeled fibers were virtually absent from the ST while they were prominent in the lateral hypothalamic area (Fig. 1g, four out of four mice tested). These results suggest that the STB^mNAc^ contains a population of mNAc-projecting fibers originating from the pBLA.

The dense fiber organization of the STB^mNAc^ provides an anatomical basis for the selective stimulation of the afferents to the mNAc. To test whether optogenetic stimulation of the STB^mNAc^ engages its source neurons, we expressed channelrhodopsin-2 (ChR2) in mNAc-projecting neurons using a retrograde AAV carrying a ChR2-tdTomato gene and delivered light to the ST (Fig. 1h). Unilateral stimulation (10 ms pulses delivered at 40 Hz) of the ChR2-expressing STB^mNAc^ produced robust c-Fos immunoreactivity in the pBLA on the stimulated side, with markedly more c-Fos-positive nuclei than in the contralateral pBLA (Fig. 1i, j; Wilcoxon signed-rank tests; *P =* 0.40, *n =* 6 for control mice (Cont); *P =* 0.031, *n =* 6 for ChR mice). The difference in c-Fos immunoreactivity between illuminated and intact hemispheres was significantly greater in ChR2-expressing mice than in control mice (Fig. 1j, inset; Wilcoxon rank sum test, *P <* 0.01, *n =* 6 per group). These findings suggest that illumination of ChR2-expressing STB^mNAc^ effectively activates a population of pBLA neurons ipsilateral to the illuminated hemisphere.

### Differential effects of fiber bundle stimulation in the two-port choice ICSS nose-poking task

Next, we examined whether and how selective engagement of the STB^mNAc^ pathway influenced reward-related behavior. To this end, we examined the behavioral effects of optogenetic stimulation in a group of mice in which ChR2-expressing axons projecting to the mNAc were photostimulated at the ST (STB^mNAc^ mice) and compared with those in another group of mice in which ChR2-expressing axons originating from the lNAc were photostimulated at the MFB (MFB^lNAc^ mice) (Fig. 2a). We first assessed the effect on ICSS behavior in a two-port choice nose-poking task (Fig. 2b, c). In this task, photostimulation (10-ms pulses at 40 Hz) was initiated upon nose-poking into one of two ports and terminated upon withdrawal. Each port was located in a separate compartment, and mice equipped with a wireless receiver connected to the implanted LED cannula (see Materials and Methods) could freely move between the compartments (Fig. 2c). Both STB^mNAc^ and MFB^lNAc^ stimulation increased cumulative nose-poking time at the stimulation-associated (active) port compared to that at the stimulation-unassociated (inactive) port in each session (Fig. 2d). Total nose-poking time in four sessions was significantly longer at the active port relative to the inactive port in both stimulated mouse groups (Fig. 2e; paired t tests; *P <* 0.001, *n =* 8 for STB^mNAc^ mice (STB-ChR); *P <* 0.01, *n =* 8 for MFB^lNAc^ mice (MFB-ChR)), whereas neither control group showed a significant difference between ports (paired t tests; *P =* 0.33, *n =* 6 for STB^mNAc^-tdTomato control mice (STB-Cont); *P =* 0.44, *n =* 6 for MFB^lNAc^-mCherry control mice (MFB-Cont)). However, plotting the duration of each nose-poking event over the course of a session revealed differences in event frequency and the distribution of event durations between STB^mNAc^ and MFB^lNAc^ mice (Fig. 2f). Statistically, the total number of nose-poking events across four sessions (see Materials and Methods) was substantially higher at the active port than at the inactive port in STB^mNAc^ mice (Fig. 2g, left; paired t test, *P <* 0.001, *n =* 8), whereas the difference was much smaller, although significant, in MFB^lNAc^ mice (paired t test, *P <* 0.001, *n =* 8). Among mouse groups, the difference between the ports was significantly larger in STB^mNAc^ mice (*n =* 8) compared to STB^mNAc^ control mice (*n =* 6) and to MFB^lNAc^ mice (*n =* 8) (Fig. 2g, right; one-way ANOVA, *P <* 0.001; Tukey’s HSD post hoc tests, *P <* 0.001 for both comparisons), but no significant difference was observed between MFB^lNAc^ mice and MFB^lNAc^ control mice (*n =* 6) (*P =* 0.075). Conversely, the mean duration of nose-poking events at the active port was only slightly increased in STB^mNAc^ mice but substantially prolonged in MFB^lNAc^ mice (Fig. 2h, left; paired t tests; *P =* 0.014, *n =* 8 for STB^mNAc^ mice; *P <* 0.01, *n =* 8 for MFB^lNAc^ mice). The difference between the ports was significantly larger in MFB^lNAc^ mice (*n =* 8) compared to MFB^lNAc^ control mice (*n =* 6) and to STB^mNAc^ mice (*n =* 8) (Fig. 2h, right; one-way ANOVA, *P <* 0.001; Tukey’s HSD post hoc tests, *P <* 0.01 for STB-ChR vs. MFB-ChR, *P <* 0.001 for MFB-Cont vs. MFB-ChR), but was not significant between STB^mNAc^ mice and STB^mNAc^ control mice (*n =* 6) (*P =* 0.27). Notably, long-duration events (> 5 s) were remarkably more frequent at the active port than at the inactive port in MFB^lNAc^ mice (Fig. 2i; paired t test, *P <* 0.001, *n* = 8), but no significant difference was observed in STB^mNAc^ mice (Wilcoxon signed-rank test, *P =* 0.20, *n* = 8), or in either control group (paired t tests; *P =* 0.65, *n* = 6 for STB-Cont; *P =* 0.90, *n* = 6 for MFB-Cont). These results indicate that STB^mNAc^ stimulation preferentially increased the frequency of nose-poking events, with only a modest effect on their duration. In contrast, MFB^lNAc^ stimulation preferentially prolonged individual nose-poking events, without a significant increase in event frequency relative to the MFB control group.

**Figure 2.**
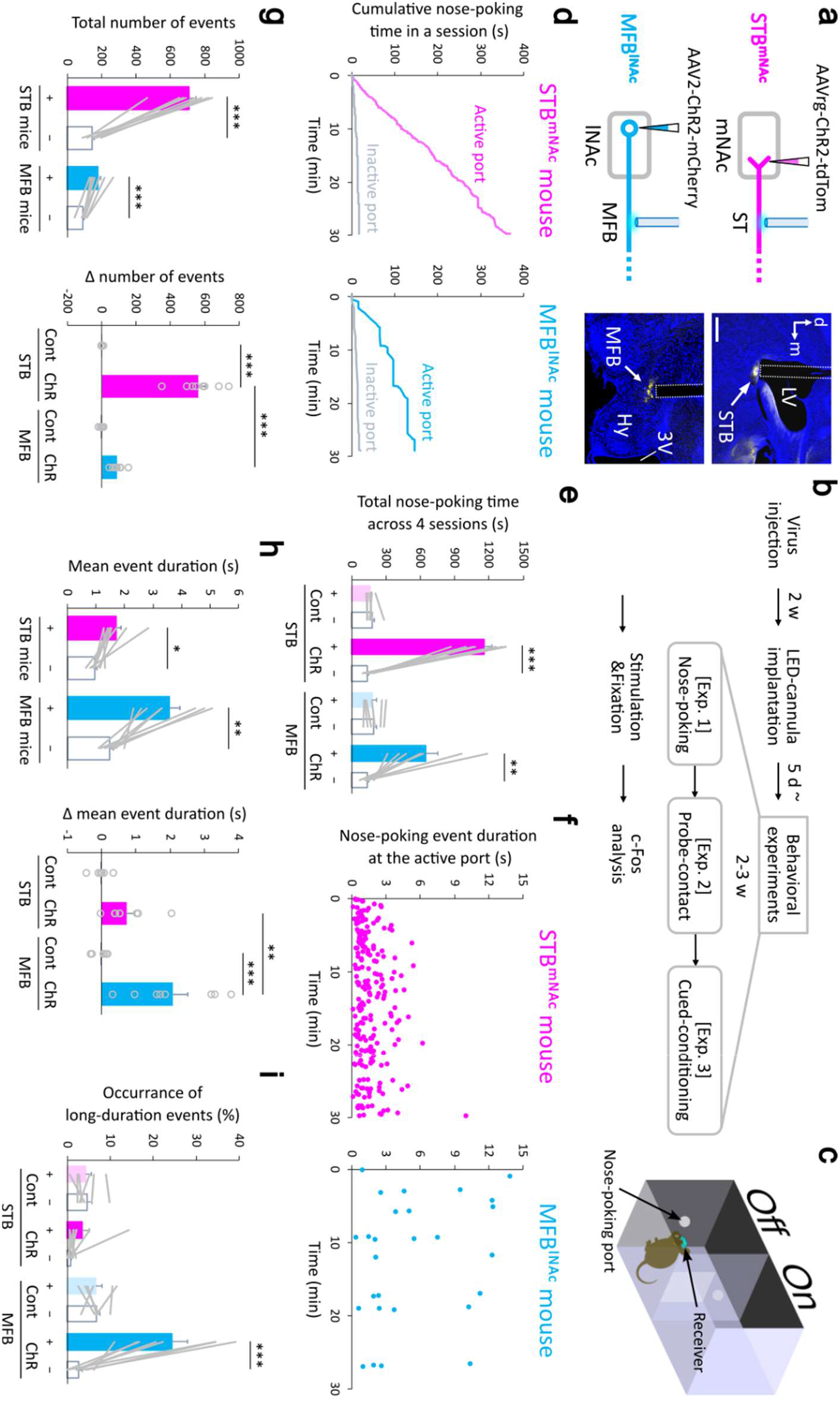
Distinct reward-related effects of fiber bundle stimulation in a two-port choice nose-poking task. **a** Left: Schematic representation of the two types of ChR2-expressing mice with fiber bundle photostimulation. Top, ChR2-expressing mNAc afferents are photostimulated at the ST (STB^mNAc^ mice). Bottom, ChR2-expressing lNAc efferents are photostimulated at the MFB (MFB^lNAc^ mice). Right: Representative coronal sections with optical fiber tracks (dotted lines) in an STB^mNAc^ (top) and an MFB^lNAc^ (bottom) mouse. Scale bar: 0.5 mm. Abbreviations: 3V, third ventricle; d, dorsal; Hy, hypothalamus; m, medial. **b** Schematic of the experimental schedule in this optogenetic study. Two weeks after virus injection, an LED cannula was implanted into the brain (a), and the mouse was maintained for 5 d or more. The mouse was then subjected to a series of behavioral experiments composed of a nose-poking task [Exp. 1], a probe-contact task [Exp. 2], and cued conditioning [Exp. 3], which were separated for at least a day, and all the experiments were completed in 2–3 weeks. More than 1 d after the behavioral experiments, mice were photostimulated and perfused for c-Fos analysis (see Materials and Methods). **c** Schematic representation of the two-port choice ICSS nose-poking task. Mice with a wireless receiver (blue) connected to the implanted LED cannula could freely move across the two compartments and nose-poke through a small window on the wall (nose-poking port) for each compartment; either one of the two nose-poking ports was paired with photostimulation (active port, On), whereas the other was unpaired (inactive port, Off). **d** Cumulative nose-poking time in a 30-min session for an STB^mNAc^ (red) and an MFB^lNAc^ (blue) mouse. Cumulative nose-poking time was greater at the active port (red or blue) than at the inactive port (gray) in both STB^mNAc^ and MFB^lNAc^ mice. **e** Comparison of total nose-poking time in four daily sessions between the active (+) and inactive (−) ports. ChR2-expressing mice (ChR), but not control mice (Cont), exhibited significantly longer nose-poking time for the active port than for the inactive port (paired t tests; *t*(5) = −1.07, *P =* 0.33, *n =* 6 for STB-Cont; *t*(7) = 14.34, *P <* 0.001, *n =* 8 for STB-ChR; *t*(5) = −0.85, *P =* 0.44, *n =* 6 for MFB-Cont; *t*(7) = 4.75, *P <* 0.01, *n =* 8 for MFB-ChR). **f** Nose-poking event duration plotted against time in a session for the active port, for an STB^mNAc^ mouse (left) and an MFB^lNAc^ mouse (right). **g** Comparison of the total number of nose-poking events across four daily sessions. Left, both ChR2-expressing STB^mNAc^ mice (red) and MFB^lNAc^ mice (blue) exhibited a significantly larger number of nose-poking events to the active port than to the inactive port (paired t tests; *t*(7) = 13.41, *P* < 0.001, *n* = 8 for STB^mNAc^ mice; *t*(7) = 7.19, *P* < 0.001, *n* = 8 for MFB^lNAc^ mice). Right, the number of nose-poking events for the active port relative to that for the inactive port. One-way ANOVA revealed a significant group effect (*F*(3,24) = 122.5, *P <* 0.001). Tukey’s HSD post hoc test revealed significant differences between STB-Cont (n = 6) and STB-ChR (n = 8) (*P <* 0.001), and between STB-ChR and MFB-ChR (n = 8) (*P <* 0.001). No significant difference was observed between MFB-Cont (n = 6) and MFB-ChR (*P =* 0.075). **h** Comparison of mean nose-poking event durations across four daily sessions. Left, both ChR2-expressing STB^mNAc^ mice (red) and MFB^lNAc^ mice (blue) exhibited significantly longer event durations for the active port than for the inactive port (paired t tests; *t*(7) = 3.28, *P* = 0.014, *n* = 8 for STB^mNAc^ mice; *t*(7) = 4.83, *P* < 0.01, *n* = 8 for MFB^lNAc^ mice). Right, mean nose-poking event duration for the active port relative to that for the inactive port. One-way ANOVA revealed a significant group effect (*F*(3,24) = 12.4, *P <* 0.001). Tukey’s HSD post hoc test revealed significant differences between STB-ChR (*n* = 8) and MFB-ChR (*n* = 8) (*P <* 0.01), and between MFB-Cont (*n* = 6) and MFB-ChR (*P <* 0.001). No significant difference was detected between STB-Cont (*n* = 6) and STB-ChR (*P =* 0.27). **i** Comparison of the occurrence of long-duration nose-poking events across four sessions between the active (+) and inactive (−) ports. Only mice with ChR2-expressing lNAc axons illuminated at the MFB exhibited significantly higher occurrence for the active port than for the inactive port (paired t tests: *t*(5) = −0.49, *P =* 0.65, *n =* 6 for STB-Cont; *t*(5) = −0.13, *P =* 0.90, *n =* 6 for MFB-Cont; *t*(7) = 5.75, *P <* 0.001, *n =* 8 for MFB-ChR; Wilcoxon signed-rank test: *V* = 28, *P =* 0.20, *n =* 8 for STB-ChR). Gray lines or circles and bars in e and g-i indicate individual mice and means ± SEM, respectively. \**P <* 0.05, \*\**P <* 0.01, \*\*\**P <* 0.001.

### Differential effects on ongoing seeking behavior

If seeking behavior is driven by reward anticipation, the occurrence and duration of subsequent behavior may depend on whether the anticipated reward is acquired. In contrast, if seeking behavior is sustained by stimulation itself, its persistence may depend on the duration of stimulation. To examine the effect of concurrent stimulation on ongoing seeking behavior, we performed another behavioral task in which mice received lasting photostimulation upon probe contact with variable stimulation durations; the duration was randomly selected from a set of four values (3, 2, 1, or 0 s [no stimulation]) in a trial-by-trial manner (Fig. 3a, also see Materials and Methods). STB^mNAc^ mice exhibited brief probe-contact behavior during stimulation, with contact time remaining similar across stimulation durations (Fig. 3b, left). Interestingly, seven of eight mice showed longer contact behavior under the no-stimulation condition than under stimulation conditions (Fig. 3b, left). Although the Friedman test revealed a significant effect of stimulation condition (*P =* 0.034, *n* = 8), none of the post hoc pairwise comparisons reached significance (Wilcoxon signed-rank test with Holm correction, all *P* ≥ 0.094). However, when the three stimulation conditions were pooled, mean contact time was significantly longer when stimulation was omitted than when stimulation was delivered (Fig. 3b, inset; paired t test, *P =* 0.026, consistent with Wilcoxon signed-rank test, *P =* 0.016, *n* = 8). In contrast, mean contact time in MFB^lNAc^ mice increased with stimulation duration (Fig. 3b, right; Friedman test, *P <* 0.001, *n* = 8). The contact time was significantly longer during 3-s stimulation than during 1-s stimulation and the no-stimulation condition (Fig. 3b, right; post hoc Holm-adjusted Wilcoxon signed-rank tests, *P =* 0.047, *n* = 8 for both comparisons), although other pairwise comparisons were not statistically significant (all *P* ≥ 0.063).

**Figure 3.**
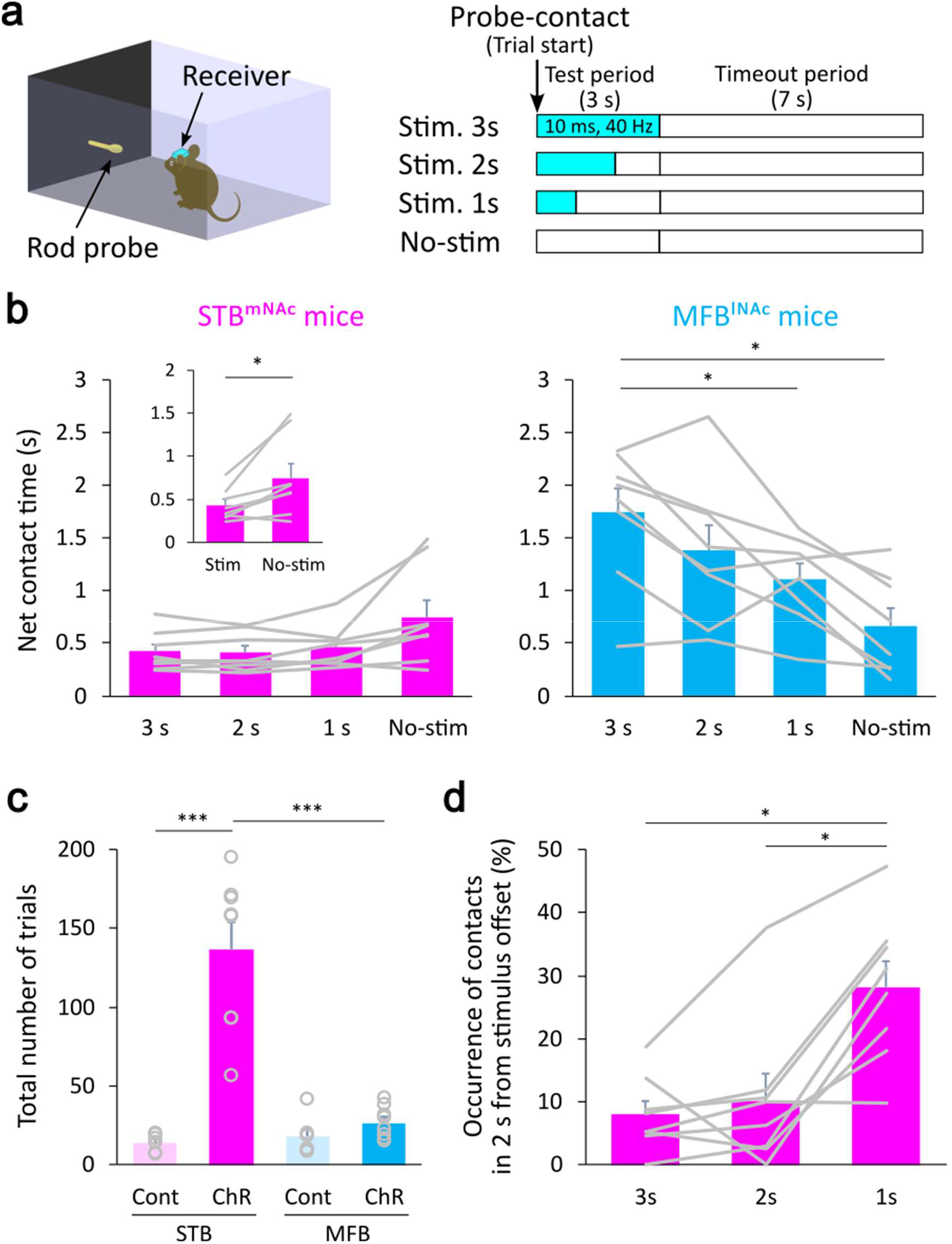
Pathway-dependent effects on ongoing seeking behavior. **a** Left: Schematic illustration of the probe-contact task. A freely moving mouse with a wireless receiver (blue) can be photostimulated upon contact with a rod probe. Right: Trial timeline and trials with variable photostimulation durations (blue). Photostimulation (10 ms, 40 Hz) was triggered by probe-contact behavior and delivered for a randomly assigned duration of 3, 2, or 1 s, irrespective of the persistence of probe contact. The stimulation was omitted in randomized contact-triggered trials (No-stim). Net probe-contact time was measured during the 3-s test period initiated by probe contact, followed by a 7-s timeout period during which photostimulation was never delivered, regardless of probe contact. **b** Net probe-contact time within the 3-s test period. Plots represent the mean net probe-contact time obtained from each stimulation condition (3-s, 2-s, and 1-s stimulation and stimulation omission [No-stim]) for each mouse. Left: STB^mNAc^ mice. A Friedman test revealed a significant difference among stimulation groups (*χ^2^*(3) = 8.70, *P =* 0.034, *n* = 8). Post hoc Wilcoxon signed-rank tests with Holm adjustment detected no significant differences between all pairs (*P =* 1.0 for [Stim. 3 s] vs. [Stim. 2 s], *P =* 0.59 for [Stim. 3 s] vs. [Stim. 1 s], *P =* 0.63 for [Stim. 2 s] vs. [Stim. 1 s], *P =* 0.094 for [Stim. 3 s] vs. [No stim], *P =* 0.094 for [Stim. 2 s] vs. [No stim], and *P =* 0.094 for [Stim. 1 s] vs. [No stim]). Inset: Comparison between the pooled stimulation conditions, including 3-s, 2-s, and 1-s stimulation, and no-stimulation condition. The mean probe-contact time was significantly longer in the no-stimulation condition than in the stimulation conditions (paired t test, *P =* 0.026; Wilcoxon signed-rank test, *P =* 0.016, *n* = 8). Right: MFB^lNAc^ mice. A Friedman test revealed a significant difference among stimulation groups (*χ^2^*(3) = 16.4, *P <* 0.001, *n* = 8). Post hoc Holm-adjusted Wilcoxon signed-rank tests revealed significant differences between [Stim. 3 s] and [No-stim] (*P =* 0.047), and between [Stim. 3 s] and [Stim. 1 s] (*P =* 0.047). No significant differences were detected between [Stim. 3 s] and [Stim. 2 s] (*P =* 0.11), [Stim. 2 s] and [Stim. 1 s] (*P =* 0.20), [Stim. 2 s] and [No-stim] (*P =* 0.063), or [Stim. 1 s] and [No-stim] (*P =* 0.070). **c** Total number of probe-contact trials. One-way ANOVA revealed a significant group effect (*F*(3,24) = 33.8, *P <* 0.001). Post hoc multiple comparisons using Tukey’s HSD test revealed significant differences between STB-Cont and STB-ChR (*P <* 0.001), and between STB-ChR and MFB-ChR (*P <* 0.001). No significant difference was found between MFB-Cont and MFB-ChR (*P =* 0.93). **d** Comparison of the occurrence of contacts within 2 s from the photostimulation offset. A Friedman test revealed a significant effect of photostimulation duration (*χ^2^*(2) = 10.8, *P <* 0.01, *n =* 8 STB^mNAc^ mice). Post hoc Wilcoxon signed-rank tests with Holm corrections revealed significant differences between [Stim. 3 s] and [Stim. 1 s] (*P =* 0.042), and between [Stim. 2 s] and [Stim. 1 s] (*P =* 0.042), but no significant difference was found between [Stim. 3 s] and [Stim. 2 s] (*P =* 0.31). Gray circles or lines and bars in b-d indicate individual mice and means ± SEM, respectively. \**P <* 0.05, \*\*\**P <* 0.001.

Consistent with the nose-poking task results (Fig. 2g), the total number of trials per session, including trials with all stimulation durations and those with stimulation omission, was significantly higher in STB^mNAc^ mice than in either MFB^lNAc^ mice or STB-Cont mice (Fig. 3c; one-way ANOVA, *P <* 0.001; Tukey’s HSD post hoc tests, *P <* 0.001 for both comparisons), whereas no significant difference was found between MFB^lNAc^ mice and MFB-Cont mice (*P =* 0.93), indicating enhanced re-engagement of probe-contact behavior in STB^mNAc^ mice.

To further characterize the effects of stimulation duration in STB^mNAc^ mice, we assessed the occurrence of probe-contact reinitiation within 2 s after photostimulation offset. The 1-s stimulation condition resulted in a significantly higher reinitiation occurrence than longer stimulation (i.e., 2 and 3 s) (Fig. 3d; Friedman test, *P <* 0.01; post hoc Wilcoxon signed-rank tests with Holm corrections, *P =* 0.042 for [Stim. 3 s] vs. [Stim. 1 s], *P =* 0.042 for [Stim. 2 s] vs. [Stim. 1 s], and *P =* 0.31 for [Stim. 3 s] vs. [Stim. 2 s], *n =* 8 mice). These findings indicate that STB^mNAc^ stimulation consistently elicited brief probe-contact behavior irrespective of stimulation duration, while shorter stimulation was associated with a higher probability of behavioral reinitiation compared with longer stimulation durations.

### Cued operant conditioning with STB^mNAc^ stimulation in the absence of primary rewards

The facilitated reinitiation of ICSS actions observed in STB^mNAc^ mice suggests that these mice learn to anticipate the rewarding outcome of their actions. To test whether this anticipatory behavior (i.e., ICSS actions) can be elicited by a cue, we performed cued operant conditioning in the absence of primary rewards. In this paradigm, photostimulation was delivered upon probe contact only during a 10-s auditory conditioned stimulus (CS) (Fig. 4a), without providing food or water rewards and without prior deprivation. Over the course of five daily sessions, STB^mNAc^ mice, all of which met the habituation criterion (6 of 6 mice; see Materials and Methods), showed progressively greater selectivity of probe-contact behavior for the CS period (Fig. 4b, red; repeated-measures ANOVA, *P <* 0.001, *n =* 6) and a reduced response latency from CS onset (Fig. 4b, gray; repeated-measures ANOVA, *P <* 0.001, *n =* 6). The selectivity index (see Materials and Methods) increased significantly relative to Day 1 on Days 3–5 (post hoc Dunnett tests, *P =* 0.028 for Day 3, *P <* 0.01 for Day 4 and Day 5), whereas response latency was significantly reduced on Days 2, 3, and 5 (post hoc Dunnett tests, *P =* 0.022 for Day 2, *P <* 0.01 for Day 3 and Day 5). STB^mNAc^ mice also showed an increase in the number of trials with probe contact during the CS across the 4 daily sessions (Fig. 4c; repeated-measures ANOVA, *P <* 0.001; post hoc Dunnett tests comparing each day with Day 1, *P =* 0.044 for Day 2, *P =* 0.043 for Day 3, and *P =* 0.046 for Day 4; *n =* 6). Conversely, the number of trials with probe contact during the pre-CS decreased (repeated-measures ANOVA, *P <* 0.001; post hoc Dunnett test comparing Day 4 with Day 1, *P =* 0.020; *n =* 6). In contrast, MFB^lNAc^ mice that met the habituation criterion (7 of 8 mice) showed no significant increase in cue-evoked probe-contact behavior across training sessions (Fig. 4c; CS: repeated-measures ANOVA, *P =* 0.13; pre-CS: repeated-measures ANOVA, *P =* 0.14; *n =* 7). These results demonstrate that STB^mNAc^ stimulation, but not MFB^lNAc^ stimulation, supports the acquisition of cue-dependent probe-contact behavior in the absence of primary rewards.

**Figure 4.**
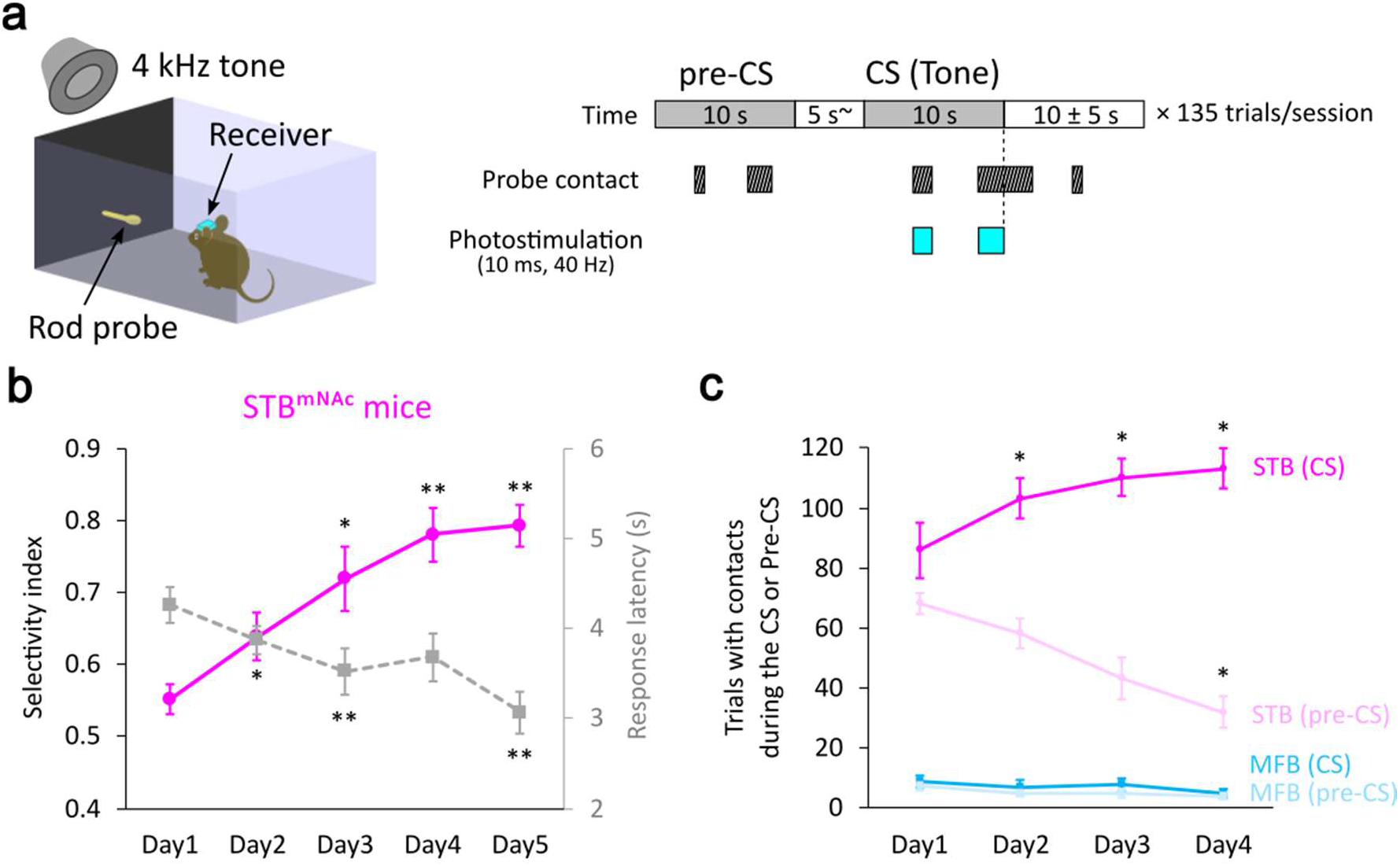
STB^mNAc^ stimulation, but not MFB^lNAc^ stimulation, can substitute for a reward in cued operant conditioning. **a** Schematic illustrating the cued operant conditioning task (left) and trial timeline (right). A freely moving mouse with a wireless receiver can acquire photostimulation upon contact with the rod probe only during the conditioned stimulus (CS; 4 kHz tone pulses). A trial was composed of a 10-s period before the CS (pre-CS period), a 10-s CS period, and a 10 ± 5-s post-CS period. The pre-CS and CS periods were separated by a 5-s timeout, during which mice were punished with an additional 5-s timeout if they contacted the probe. A daily training session contains 135 trials, and mice were trained for 5 (STB^mNAc^ mice) or 4 (MFB^lNAc^ mice) consecutive days. **b** An increase in the selectivity index (red, see Materials and Methods) and a decrease in behavioral response latency (gray) over conditioning days in STB^mNAc^ mice. A repeated-measures ANOVA revealed a significant difference among days (*F*(4, 20) = 22.9, *P <* 0.001 for the selectivity index; *F*(4, 20) = 11.9, *P <* 0.001 for the response latency, *n =* 6 mice). Post hoc Dunnett tests revealed significant differences between Day 1 and Day 3 (*P =* 0.028), Day 4 (*P <* 0.01), and Day 5 (*P <* 0.01), but not Day 2 (*P =* 0.053), for the selectivity index, and between Day 1 and Day 2 (*P =* 0.022), Day 3 (*P <* 0.01), and Day 5 (*P <* 0.01), but not Day 4 (*P =* 0.13), for the response latency. **c** Total number of trials with probe contacts during the CS or pre-CS across conditioning days. A repeated-measures ANOVA revealed a significant difference among days in STB^mNAc^ mice (*F*(3, 15) = 9.34, *P <* 0.001 for CS (red); *F*(3, 15) = 11.8, *P <* 0.001 for pre-CS (light pink); *n =* 6), but not in MFB^lNAc^ mice (*F*(3, 18) = 2.13, *P =* 0.13 for CS (blue); *F*(3, 18) = 2.09, *P =* 0.14 for pre-CS (light blue); *n =* 7). In STB^mNAc^ mice, post hoc Dunnett tests revealed significant differences between Day 1 and Day 2 (*P =* 0.044), Day 3 (*P =* 0.043), and Day 4 (*P =* 0.046) for CS, and between Day 1 and Day 4 (*P =* 0.020), but not Day 2 (*P =* 0.31) and Day 3 (*P =* 0.11) for pre-CS. The plots and error bars represent means ± SEM. \**P <* 0.05, \*\**P <* 0.01.

## Discussion

### Distinct components of reward-related behavior

Reward-related behavior comprises multiple processes, including the initiation and persistence of reward-seeking actions, their termination, and the subsequent reinitiation of actions. Although these processes are functionally related, they may be differentially regulated by distinct neural pathways. In the present study, both STB^mNAc^ and MFB^lNAc^ stimulation promoted ICSS behavior, but their effects differed markedly in their temporal relationship to stimulation: STB^mNAc^ stimulation promoted the reinitiation of actions during periods when stimulation was absent, whereas MFB^lNAc^ stimulation predominantly prolonged ongoing behavior during stimulation. This dissociation suggests that the reinitiation and persistence of motivated behavior are differentially regulated by distinct neural substrates. The reinitiation of ICSS actions in STB^mNAc^ mice further suggests that these actions are not directly elicited by concurrent stimulation but may instead reflect reward anticipation based on previously acquired incentive value. Consistent with this interpretation, STB^mNAc^ stimulation supported the acquisition of preference for the stimulation-associated port and probe, as well as cue-dependent operant behavior.

The duration of STB^mNAc^ stimulation also influenced the temporal dynamics of seeking behavior. Probe-contact time was similar across stimulation durations, whereas contact time in the stimulation-omission condition was longer on average. Furthermore, reinitiation of probe-contact behavior within 2 s after stimulation offset was more likely following 1-s stimulation than following 2- or 3-s stimulation. STB^mNAc^ mice also initiated substantially more probe-contact trials than control or MFB^lNAc^ mice, indicating enhanced repeated engagement in seeking behavior. Taken together, these findings raise the possibility that the acquisition of reward during STB^mNAc^ stimulation transiently terminates ongoing seeking behavior, whereas the absence or insufficiency of expected rewarding stimulation may favor subsequent behavioral re-engagement. Thus, both the duration and presence or absence of STB^mNAc^ stimulation may influence the termination and subsequent reinitiation of reward-seeking behavior, potentially reflecting distinct temporal components of reward acquisition and motivational processes.

Importantly, the acquisition of preference and cue-dependent operant behavior in the absence of primary rewards suggests that STB^mNAc^ stimulation confers a positive incentive value. However, the increase in nose-poking event duration in STB^mNAc^ mice was not significantly greater than that in control mice. It is unlikely that the stimulation directly suppressed motor output because the nose-poking behavior at the active port lasted for approximately 1.7 s on average (Fig. 2h). It is therefore possible that the processes underlying incentive value and the persistence of motivated behavior are functionally dissociable. This apparent dissociation also raises the possibility that reward-related processes associated with STB^mNAc^ stimulation contribute to the regulation of ongoing behavior. Whether the termination of reward-seeking behavior during STB^mNAc^ stimulation is caused by a process analogous to reward consumption or transient satiety remains to be determined.

Although MFB^lNAc^ stimulation maintained ongoing behavior, the cue-dependent operant response was not significantly increased by MFB^lNAc^ stimulation across training sessions under the conditions of the present paradigm. This finding suggests that the persistence of motivational drive is not necessarily sufficient to establish cue-dependent anticipatory behavior. Rather, the acquisition of such behavior may require learning of reward incentive value that enables subsequent reward anticipation.

### STB^mNAc^ stimulation as a tool for isolating reward-dependent processes

Primary rewards engage sensory, motivational, homeostatic, and affective processes simultaneously, making it difficult to isolate the neural processes underlying reward incentive value. In the present study, STB^mNAc^ stimulation promoted re-engagement of seeking behavior in the absence of prior deprivation, indicating that its behavioral effects do not require an experimentally elevated homeostatic drive, as is often required to promote robust seeking behavior in laboratory settings (*36–38*). Moreover, because STB^mNAc^ stimulation supported reward-dependent behavior without primary rewards, it may provide access to neural processes underlying reward-related behavior independently of peripheral sensory transduction. Thus, STB^mNAc^ stimulation may offer an experimental approach for investigating reward-related processes while reducing the confounding influences of sensory modality and homeostatic regulation (*3*). Nevertheless, whether the affective state elicited by STB^mNAc^ stimulation shares specific features with that induced by natural rewards remains to be established.

### Possible neural circuits underlying the effects of fiber bundle stimulation

What neural circuits mediate the behavioral effects of STB^mNAc^ and MFB^lNAc^ stimulation? A previous study implicated the mNAc-VTA circuit in the suppression of dopaminergic activity in the VTA (*16*), raising the possibility that STB^mNAc^ stimulation inhibits the persistence of reward-seeking behavior through this pathway. However, because activation of mNAc neurons did not significantly affect preference acquisition in previous studies (*16, 17, 39*), reward incentive value conferred by STB^mNAc^ stimulation may involve additional pathways. A genetically defined population of pBLA neurons has been implicated in positive valence, although its projections to the NAc were not predominantly distributed to the medial portion (*7, 20*). Based on our anatomical findings (Fig. 1e, f), STB^mNAc^ stimulation in the present study may preferentially recruit an mNAc-projecting subpopulation of

### pBLA neurons associated with positive valence

In the MFB^lNAc^ stimulation experiments, photostimulation was restricted to lNAc-derived axons coursing through the MFB, a heterogeneous fiber bundle containing both ascending and descending axons (*35, 40, 41*). Among the descending fibers, our MFB^lNAc^ stimulation may have preferentially activated the direct lNAc-VTA pathway over the indirect pathway via the ventral pallidum (*42*). This possibility is compatible with a previous study demonstrating that photostimulation of lNAc axon terminals in the VTA reinforced ICSS (*16*). Given the involvement of the lNAc in promoting dopaminergic activity (*16*), our MFB^lNAc^ stimulation may maintain motivational drive by recruiting VTA dopaminergic neurons. Nevertheless, stimulation of STB^mNAc^ and MFB^lNAc^ fibers may recruit multiple downstream pathways and potentially engage collateral circuits through antidromic activation (*35*). Future studies using projection-specific circuit manipulations will be necessary to identify the pathways responsible for the reinitiation and persistence of motivated behavior, as well as those involved in the generation of reward incentive value.

## Conclusion

In conclusion, our findings reveal a functional dissociation between STB^mNAc^ and MFB^lNAc^ stimulation in reward-related behavior. Whereas MFB^lNAc^ stimulation predominantly sustains ongoing actions, STB^mNAc^ stimulation promotes the reinitiation of seeking actions and supports reward-dependent learning in the absence of primary rewards. These findings suggest that reward-related behavior is not governed by a unitary motivational system but involves anatomically and functionally distinct neural components that regulate the persistence of motivated behavior, the acquisition of reward incentive value, and the subsequent reinitiation of actions. By providing an experimentally accessible means of investigating reward-related processes with reduced dependence on sensory modality and homeostatic drive, STB^mNAc^ stimulation offers a framework for studying the neural and physiological substrates of reward processing. More broadly, the functional dissociation identified here may be helpful for the development of therapeutic brain stimulation strategies targeting distinct components of reward-related dysfunction (*43*).

## Materials and Methods

### Animals

All experiments were performed in male C57BL/6J (B6) mice (Japan SLC, Shizuoka, Japan). The mice were 6–10 weeks old and weighed 20–25 g at the time of virus injection (see below). Mice were group-housed before virus injection and individually housed and maintained thereafter in a breeding room on a 12 h light-dark cycle with food and water available ad libitum. All experiments were approved by the Committee for Animal Experiments of Kumamoto University and were performed in accordance with the Guidelines for Use of Animals in Experiments of Kumamoto University.

### Virus injections

Mice were deeply anesthetized with a mixture of ketamine (Ketamine injection, Fujita Pharmaceutical, Tokyo, Japan) and xylazine (Selactar, Bayer Yakuhin, Tokyo, Japan) (80 mg/kg and 8 mg/kg, respectively) and head-fixed on a stereotaxic frame (SR-5M-HT, Narishige, Tokyo, Japan). After partial hair removal and incision of the skin along the midline of the head to allow skull exposure, small craniotomies (< 1 mm square each) were made over the locations of the virus injections. Virus injections were performed using air pressure (0.1 µl per 2–3 min) with a custom-made syringe pump connected to a glass capillary needle (Calibrated Pipets #2-000-001-90, 30-40 µm tip diameter, Drummond Scientific Company, PA, USA). For retrograde labeling of neurons projecting to the NAc, ∼0.1 µL of an AAV vector, retrograde AAV-CAG-tdTomato (pAAV-CAG-tdTomato (codon diversified) was a gift from Edward Boyden, Addgene viral prep # 59462-AAVrg; http://n2t.net/addgene:59462; RRID: Addgene_59462, titer: 1 × 10^13^ genome copies (GC)/mL, Addgene, MA, USA) was injected into the mNAc (1.1 mm anterior, 0.5 mm lateral, and 4.4 mm ventral to bregma) of the left hemisphere. For selective labeling of mNAc-projecting neurons in the pBLA and mVTA, ∼0.2 µL of a retrograde AAV-hSyn-Cre (pENN.AAV.hSyn.Cre.WPRE.hGH was a gift from James M. Wilson, Addgene viral prep # 105553-AAVrg; http://n2t.net/addgene:105553; RRID:Addgene_105553, titer: 1 × 10^13^ GC/mL), ∼0.15 µL of an AAV2-CAG-FLEX-GFP (titer: 4 × 10^12^ GC/mL, University of North Carolina (UNC) Vector Core, NC, USA), and an AAV2-CAG-FLEX-tdTomato (pAAV-FLEX-tdTomato was a gift from Edward Boyden, Addgene viral prep # 28306-AAV2; http://n2t.net/addgene:28306; RRID:Addgene_28306, titer: 9 × 10^12^ GC/mL) were injected into the mNAc, pBLA (2.1 mm posterior, 3.1 mm lateral, 5.0 mm ventral to bregma), and mVTA (3.1 mm posterior, 0.4 mm lateral, and 4.7 mm ventral to bregma) of the left hemisphere, respectively. For optogenetic experiments (see below), either a retrograde AAV-CAG-hChR2(H134R)-tdTomato (pAAV-CAG-hChR2-H134R-tdTomato was a gift from Karel Svoboda, Addgene viral prep # 28017-AAVrg; http://n2t.net/addgene:28017; RRID:Addgene_28017, titer: 2 × 10^13^ GC/mL) (*44*) (for STB^mNAc^ stimulation) or a retrograde AAV-CAG-tdTomato (for STB^mNAc^ control) was injected into the mNAc, and either an AAV2-CAG-hChR2(H134R)-mCherry (pAAV.CAG.hChR2(H134R)-mCherry.WPRE.SV40 was a gift from Karl Deisseroth, Addgene viral prep # 100054-AAV2; http://n2t.net/addgene:100054; RRID:Addgene_100054, titer: 3 × 10^12^ GC/mL) (for MFB^lNAc^ stimulation) or an AAV2-hSyn-mCherry (titer: 5 × 10^12^ GC/mL, UNC Vector Core) (for MFB^lNAc^ control) was injected into the lNAc (1.0 mm anterior, 1.8 mm lateral, and 4.9 mm ventral to bregma) of the left hemisphere. After the injections, the parietal skin was sutured and glued with elastic adhesive. The mice were then injected subcutaneously with buprenorphine (0.1 mg/kg body weight, Lepetan, Otsuka Pharmaceutical, Tokyo) and enrofloxacin (5 mg/kg body weight, Baytril, Bayer Yakuhin, Tokyo) for postoperative analgesia and protection against bacterial infections, respectively. The mice were maintained for one and two weeks until sacrifice for retrograde tracing and axon tracing, respectively, and for two weeks for LED cannula implantation (see below).

### LED cannula implantation

An LED cannula was fabricated by gluing one end of a 200-µm-diameter optical fiber (FG200UCC, Thorlabs, 5-7 mm in length) to the center of a blue surface-mount LED (LB G6SP, ams OSRAM, Munich, Germany). The three electrodes of the LED were each soldered to separate connectors to facilitate connection with a wireless receiver (see below). The light intensity of the LED cannula was measured using an optical power meter (LPM-100, Bio Research Center, Nagoya, Japan), and LED cannulae with an output of 4.5 ± 1.5 mW were used. For LED cannula implantation, craniotomy was performed in the same manner as for virus injections, and the optical fiber was then inserted stereotaxically into the stimulation sites. The coordinates for the fiber tips were 1.0 mm posterior, 2.0 mm lateral, and 2.7 mm ventral (for ST illumination) and 1.3 mm posterior, 1.4 mm lateral, and 4.6 mm ventral (for MFB illumination) to bregma. A screw was embedded in the frontal part of the skull, and the LED cannula and screw were covered with resin cement (Super-Bond, Sun Medical, Shiga, Japan) and elastic adhesive. After subcutaneous injections of buprenorphine and enrofloxacin (see above), the mice were individually housed and maintained in a breeding room for at least 5 days after LED implantation until behavioral experiments.

### Optogenetics

Optogenetic stimulation was controlled using a wireless system (Teleopto, Bio Research Center). At the start of each session of all behavioral experiments with optogenetics, a 1.4-g wireless receiver was connected to the animal’s connector under light inhalation anesthesia with isoflurane (4% for induction and 2% for maintenance). The receiver was fully charged prior to each session. The receiver was controlled by a remote controller through an infrared emitter. To confirm the efficacy of the optogenetic stimulation, c-Fos analysis (see below) was performed after the end of all behavioral experiments.

### Behavioral experiments

#### Two-port-choice ICSS nose-poking task

Four daily sessions (30 min from the first nose-poking event for each session) were conducted for consecutive days. After connecting a receiver (see above), mice were maintained in a transfer cage for more than 10 min to fully recover from anesthesia. The mice were then placed in a custom-made nose-poking chamber, which was placed in a sound-attenuated box. The chamber consisted of two compartments connected by a short passage through which mice could pass freely. Each compartment (130 × 130 × 100 mm) had a small window on the wall (∼10 mm in diameter, 2.5 cm above the floor) to which mice spontaneously engaged in nose-poking as an exploratory behavior. Either one of the windows (referred to as the active port) was paired with LED illumination, while the other one (referred to as the inactive port) was unpaired. LED illumination was initiated and terminated upon the onset and offset of nose-poking at the active port, respectively, and consisted of 10-ms light pulses delivered at 40 Hz. To minimize the influence of unexpected selection bias, the active and inactive ports were switched every session (the right window was assigned to the active port for the first and third sessions). Nose-poking behavior was detected using an infrared sensor module (Main chip LM393; Optocoupler H2010, DAOKAI Co., Ltd., Foshan, China) mounted on the outside of each port. Data were collected using an Arduino microcomputer (Arduino UNO R3, Arduino, https://www.arduino.cc/). Nose-poking lasting more than 0.1 s was detected as a behavioral event. Successive behavioral bouts that were intercalated by a non-nose-poking period of less than 0.75 s were regarded as a single continuous event. Cumulative nose-poking time, total number of events, and mean duration of events were analyzed for each session.

#### Probe-contact task with variable stimulation duration

After recovering from anesthesia for the connection of a wireless receiver (see above), the mice were placed in a chamber equipped with a metal rod probe on a wall (∼ 4 cm above the floor) and an aluminum sheet on the floor. Probe contact was detected using a capacitive touch sensor (LK-CB6, EK JAPAN, Fukuoka, Japan) connected to the rod probe. Contacting the probe triggered 10 ms pulses of LED illumination at 40 Hz for randomized durations of 3, 2, 1, and 0 s (i.e., stimulation omission) on a trial-by-trial basis. The contact-triggered photostimulation was delivered only once per 10-s trial, regardless of additional probe-contact events during the trial. A 7-s timeout period was imposed to prevent subsequent contact-triggered stimulation within the same trial. Two daily sessions were performed for 1 h each day, with each session starting at the initial probe-contact event. The first session was assigned as a habituation session, and data were obtained during the second session. The cumulative time of probe contact behavior during the 3-s test period from the onset of initial contact for each trial, total number of trials in 1 h of the session, and number of trials with probe contacts during the 2-s period from the offset of photostimulation were quantified for each mouse. If the mice did not achieve at least three trials for each stimulation duration and stimulation omission within 1 h of the session, the session was extended until the number of trials reached three for all the stimulation conditions.

#### Cued operant conditioning with photostimulation as a substitute for the primary reinforcer

Mice were trained in a cued operant conditioning task in which optogenetic stimulation was used as a substitute for the primary reinforcer. The mice were allowed to contact a metal rod probe paired with photostimulation (10 ms pulses delivered at 40 Hz) upon contact until 20 contacts before the training session (only on the first day). A single contact was counted as a contact behavior lasting within 0.5 s during the habituation period. Only mice that had achieved 20 contacts in 1 h of the habituation period were permitted to move on to the first session of training immediately thereafter. Of the mice included in the analysis, all six STB^mNAc^ mice, none of the six STB^mNAc^ control mice, seven of eight MFB^lNAc^ mice, and one of six MFB^lNAc^ control mice satisfied this habituation criterion. Note that all mice had completed both the nose-poking and probe-contact tasks by the first day of the conditioning task. STB^mNAc^ mice underwent five consecutive daily training sessions, whereas MFB^lNAc^ mice underwent four consecutive daily sessions, with 135 trials per session. Each trial consisted of a 10-s conditioned stimulus (CS) period associated with a sound cue (250 ms 4 kHz tone pulses presented at 2 Hz, 65 dB SPL), during which a train of 470 nm LED illumination (10 ms pulse at 40 Hz) was delivered upon the mouse’s contact with a rod probe (see above), and a 10-s pre-CS period that was separated by a 5-s period from the CS period. Probe contact was monitored continuously, and contact behavior during the 5-s period prior to the CS was punished with a 5-s timeout (from the end of the contact). Contacts occurring outside the CS period did not trigger photostimulation. Behavioral measures included the number of trials with probe contact during the CS epoch and that during the pre-CS epoch, and response latency from CS onset to the first probe contact during cue presentation. Cue presentation and photostimulation were controlled using a microcontroller-based system (Arduino UNO R3). Reinforcement learning was quantified as the selectivity index, which was calculated as X_(CS)_ / [X_(CS)_ + X_(pre-CS)_], where X_(CS)_ and X_(pre-CS)_ are the number of trials with CS and pre-CS responses, respectively. Learning was also evaluated by the reduction in response latency and an increase and decrease in the number of trials with contacts during the CS and pre-CS, respectively. Only the data obtained from mice that had started the first day of training within 3 days after the end of the probe-contact task were included in the analysis (2 of 8 STB^mNAc^ mice were excluded from the analysis due to exceeding interval days).

### Immunohistochemistry

After a series of behavioral experiments, mice were optogenetically stimulated (2 s trains of 10 ms pulses at 40 Hz delivered every 4 s) in a freely moving condition for 1 h. Thirty minutes after the end of stimulation, the mice were deeply anesthetized with a mixture of ketamine and xylazine (ketamine 100 mg/kg, xylazine 10 mg/kg), perfused with phosphate-buffered saline (PBS) and 4% paraformaldehyde (PFA) in 0.1 M phosphate buffer (PB), and brains were excised. After gradual replacement with 30% sucrose in PBS, the brains were sectioned to a thickness of 40 µm using a freezing microtome (Yamato Kohki, Saitama, Japan). For immunostaining, coronal sections were incubated in PBS containing 5% normal donkey serum and 0.3% Triton X-100 for 1 h at room temperature (RT) and then stained with primary antibodies overnight (GFP) or for 48 h (c-Fos) at 4℃ (rat anti-GFP [GF090R], 1:2000, Nacalai Tesque, Kyoto, Japan; rabbit anti-c-Fos [9F6], 1:1000, Cell Signaling Technology, MA, USA). After washing three times in PBS for 10 min, the sections were incubated with secondary antibodies for 2 h at RT (donkey anti-rat IgG Alexa Fluor^TM^ Plus-488 conjugate for GFP; donkey anti-rabbit IgG Alexa Fluor^TM^ Plus-647 or -488 conjugate for c-Fos; 1:1000 dilution for all secondary antibodies; Thermo Fisher Scientific, MA, USA). After washing three times with PBS for 10 min, the sections were mounted on glass slides and counterstained with DAPI. Subsequently, coverslips were mounted with 10% Mowiol 4–88 (Polysciences, PA, USA) in 0.1 M Tris-HCl (pH 8.0) and 25% (w/v) glycerol. Low-magnification fluorescent images of whole sections and higher-magnification images of a part of the sections were obtained using a fluorescence microscope (4 × objective lens, BZ-X710, Keyence, Osaka, Japan) and a confocal laser scanning microscope (10 × objective lens, FV3000, Olympus, Tokyo, Japan), respectively. The high-magnification images in Figures 1c, g, and i (also images used for quantification, see below) were created by a maximum intensity projection using ImageJ (NIH, MD, USA) from three confocal optical slices acquired at 2-µm intervals.

### Quantification of retrogradely labeled cells and c-Fos immunostained particles

For the quantification of retrogradely labeled tdTomato-expressing cells, 1,024 pixels (1.243 µm/pixel) square gray-scale images of the BLA (maximum intensity projection images, see above) were obtained from the virus-injected hemisphere of coronal sections. To subtract the background level using ImageJ, the mean gray values and SDs were measured for each of the four 30-pixel square areas outside the BLA, and the area with the minimal mean value was then selected to determine the mean value + 10 × SD as the background level to be subtracted from the original gray values (using the Math function). The number of labeled cells within the BLA complex (the area surrounded by the external and intermediate capsules, outlined using DAPI-stained images) was counted for each section at six levels along the anterior-posterior (A-P) axis (corresponding to 1.3 mm (AP1), 1.5 mm (AP2), 1.7 mm (AP3), 1.9 mm (AP4), 2.1 mm (AP5), and 2.3 mm (AP6) posterior to bregma of a mouse brain atlas) (*45*).

To quantify c-Fos immunoreactivity after optogenetic stimulation of the STB^mNAc^, fluorescent confocal images of the pBLA (maximum intensity projection images) were obtained from both photostimulated and unstimulated hemispheres in serial coronal sections. The A-P level was aligned to select sections for hemispheric comparisons based on the location of the amygdaloid complex relative to that of the lateral ventricle (Fig. 1i), and the region of interest (ROI) for the quantitative analysis was determined as a 200 µm square in the medial 2/3 area of the pBLA for each hemisphere. Significant c-Fos-immunoreactive particles were determined using ImageJ, as described previously (*46*), with minor modifications: to binarize the grayscale of a c-Fos immunofluorescence ROI image, a background image was created from the original grayscale image with a ‘‘rolling ball radius’’ of 50 pixels, and the mean gray value and SD of the whole ROI were measured. The original grayscale image was then thresholded using the mean + 10 × SD (for the Alexa-488 case) or the mean + 25 × SD (for the Alexa-647 case) of the background image. The binarized image was further filtered using a median filter (“radius” of 2 pixels) and segmented using the watershed function. Finally, particles ≥ 20 µm^2^ were considered significant c-Fos immunopositive particles. Overlapping particles were manually separated based on the corresponding DAPI-stained cell nuclei. The mean number of significant particles within the two ROI images obtained from the serial BLA sections was calculated for each hemisphere of the mouse.

### Statistical analyses

Statistical analyses were performed using R (version 4.5.1, the R Foundation). All data used for statistical analyses and the corresponding results are provided in Supplementary Data S1. For all pairwise comparisons other than post hoc comparisons, the Shapiro-Wilk test was used to assess the normality of the difference between groups. For the comparison of the A-P fractions of retrogradely labeled cells in the BLA, a paired t test was used to determine whether the difference was significant between A-P levels (the sum of AP1-AP3 vs. the sum of AP4-AP6, see above). For the c-Fos analysis, the Wilcoxon signed-rank test was used to test for difference between photostimulated and unstimulated hemispheres in ChR2-tdTomato-expressing STB^mNAc^ mice and tdTomato-expressing control mice. Likewise, for behavioral experiments, paired t tests or Wilcoxon signed-rank tests (only for STB^mNAc^ mice in the occurrence of long-duration events) were used to determine the significance of the differences between the active and inactive ports (nose-poking task). For multiple comparisons among mouse groups (i.e., STB^mNAc^ control, STB^mNAc^, MFB^lNAc^ control, and MFB^lNAc^ mice), a one-way ANOVA followed by Tukey’s honestly significant difference (HSD) test was used. For probe-contact time, net contact time was measured during the 3-s test period from the onset of initial probe contact (Fig. 3a) because mice tended to contact and release the probe repeatedly during the period. The mean net contact time was then calculated separately for each of the four stimulation conditions (3-s, 2-s, and 1-s stimulation and stimulation omission) for each mouse. Differences across the four conditions were assessed using the Friedman test, followed by pairwise Wilcoxon signed-rank tests with Holm correction for multiple comparisons. To compare the stimulation omission condition with stimulation overall, all contact events recorded during the three stimulation conditions (3-s, 2-s, and 1-s stimulation) were pooled for each mouse, and the mean duration was calculated across the pooled events. This value was compared with the mean contact duration in the stimulation omission condition using a paired t test. For the occurrence of probe contacts in 2 s from photostimulation offset in STB^mNAc^ mice, differences among the three related groups (3-s, 2-s, and 1-s stimulation) were assessed using the Friedman test, followed by Holm-adjusted pairwise Wilcoxon signed-rank tests. For the cued operant conditioning task, a repeated-measures ANOVA followed by Dunnett’s multiple-comparisons test was performed to compare each subsequent daily session with Day 1. Statistical significance was set at *P <* 0.05. Data are expressed as mean ± standard error of the mean unless otherwise stated.

## Acknowledgments

This work was supported by the following grants:

JSPS KAKENHI Grant Number 23K06014 (M.T.)

JSPS KAKENHI Grant Number 22K06433 and 25K02378 (W.J.S.)

This work was also supported by Messrs. Toshihiro Morikawa, Daisuke H. Tanaka, and 34 crowdfunding backers on the “academist” platform (https://academist-cf.com/projects/91). (M.T.)

## Author contributions to the paper

Conceptualization: M.T.; Methodology: M.T.; Investigation: M.T.; Supervision: W.J.S.; Writing–original draft: M.T.; Writing–review and editing: M.T. and W.J.S.; Funding acquisition: M.T. and W.J.S.

The authors declare no conflict of interest.

All data required to evaluate the conclusions of this study are present in the paper and/or Supplementary Materials.

We thank Dr. Edward Boyden, James M. Wilson, Karel Svoboda, and Karl Deisseroth for providing the viral vectors used in this study. We also thank Dr. Ryohei Tomioka for helpful discussions.

Paperpal and ChatGPT were used to support language editing and pre-submission checks. All final interpretations and conclusions are solely the responsibility of the authors.

## Notes

### Competing Interest Statement

The authors have declared no competing interest.

